## Supplementary material for "Coral restoration alters reef soundscapes but machine learning and manual analyses suggest different recovery rates": SI: SI.docx

Supporting Information

**Table S1 Output from a Linear Mixed Model investigating the effect of site on coral cover.** Model estimates (Emmean), standard error (SE), degrees of freedom (df) and confidence intervals are provided. The model explained a significant proportion of the variance in coral cover (Null deviance = 11663.5 on 44 degrees of freedom, Residual deviance = 4312.6 on 42 degrees of freedom, Adjusted R² = 0.619).

| site_type | Emmean | SE | df | lower.CL | upper.CL |
| --- | --- | --- | --- | --- | --- |
| degraded | 2.60 | 2.61 | 42 | -2.67 | 7.88 |
| healthy | 33.27 | 2.61 | 42 | 27.99 | 38.55 |
| outplant | 12.48 | 2.61 | 42 | 7.20 | 17.76 |

**Table S2 Post-hoc comparisons of model estimates of coral cover provided in Table S1, adjustment method was Bonferroni.** Model estimates, standard error (SE), degrees of freedom, t-ratio, and p-values are given.

| Contrast | Estimate | SE | df | t.ratio | p.value |
| --- | --- | --- | --- | --- | --- |
| degraded - healthy | -30.66 | 3.70 | 42 | -8.28 | 6.67E-10 |
| degraded - outplant | -9.87 | 3.70 | 42 | -2.66 | 0.032 |
| healthy - outplant | 20.78 | 3.70 | 42 | 5.61 | 4.21E-06 |

**Table S3 Deployment schedule for the November 2023 soundscape recording period.** Date and recorder ID (R_ID) are given for each lunar phase.

| Site | New Moon | | First Quarter | | Full Moon | | Last Quarter | | New Moon | |
| --- | --- | --- | --- | --- | --- | --- | --- | --- | --- | --- |
|  | Date | R_ID | Date | R_ID | Date | R_ID | Date | R_ID | Date | R_ID |
| outplant | 13-Nov | 1 | 19-Nov | 2 | 27-Nov | 1 | 04-Dec | 2 | x | x |
| degraded | 14-Nov | 1 | 20-Nov | 2 | 28-Nov | 1 | 05-Dec | 2 | x | x |
| healthy | x | x | 20-Nov | 1 | 28-Nov | 2 | 05-Dec | 1 | 10-Dec | 2 |

**Table S4 Description of unidentified fish sounds with comparison to previous studies.**

| Sound | Description | Type | Min frequency | Max frequency | Min duration | Num. occurrences | Paper comparison |
| --- | --- | --- | --- | --- | --- | --- | --- |
| purr | ripply purr like a cat, pulse train slow | | 300 | 600 | 1.2 | 58 | purr, Lamont 2022 |
| brrr | high pitched, like someone saying brrr | Pulse train | 300 | 800 | 0.4 | 51 | raspberry, Lamont 2022 |
| grunt | short low grunt sound | single pulse | 50 | 400 | 1 | 34 | grunt, Lamont 2022 |
| triple knocks | three triplets of high pitched knocks, two trips fast one slow | pulsed | 200 | 600 | 2.1 | 31 | knock, Lamont 2022 |
| pop | finger in cheek pop single pulse | | 300 | 1000 | 0.7 | 28 |  |
| grumble | similar to a creaky door opening but low pitch | continuous/ single pulse | 100 | 400 | 1 | 24 |  |
| robot wood pigeon | cooing but a bit robotic | continuous | 300 | 700 | 2 | 21 | croak, Lamont 2022 |
| chicken | fast plucking | pulsed | 300 | 900 | 2 | 17 |  |
| wheezing | singular exhausted inhale | single pulse | 100 | 700 | 1.3 | 15 |  |
| woodpecker like a woodpecker | | pulsed | 200 | 500 | 1 | 11 |  |
| agreement purr | falling sounds, like purr but pitch getting lower | pulsed/ single pulse? | 100 | 300 | 0.9 | 5 |  |
| cuckoo | two tone, high then low pitched | single pulse | 200 | 700 | 0.7 | 5 | whoop, Lamont 2022 |

**Table S5 Cohen’s (d) effect sizes for call rate variability by site.** Contrast refers to the sites being compared. Effect size is the Cohen’s effect size (d). SE is standard error. CL refers to the confidence level, lower and upper.

| Contrast | Effect size | SE | df | lower.CL | upper.CL |
| --- | --- | --- | --- | --- | --- |
| (degraded - healthy) | -1.86363 | 0.411736 | 39 | -2.69644 | -1.03081 |
| (degraded - outplant_south) | -1.04829 | 0.372946 | 39 | -1.80264 | -0.29393 |
| (healthy - outplant_south) | 0.815336 | 0.365408 | 39 | 0.076229 | 1.554443 |

**Table S6 Output from linear model investigating the effect of site on call rate.** The model explained a significant proportion of the variance in call rate (F(8,39) = 8.03, p < 0.001, Adjusted R² = 0.545).

| Site | Emmean | SE | df | lower.CL | upper.CL |
| --- | --- | --- | --- | --- | --- |
| degraded | 4.5 | 0.53 | 39 | 3.41 | 5.58 |
| Healthy | 8.5 | 0.53 | 39 | 7.41 | 9.58 |
| Outplant | 6.75 | 0.53 | 39 | 5.66 | 7.83 |

**Table S7 Post-hoc comparisons of model estimates of call rate provided in Table S4, adjustment method was Tukey.** Model estimates, standard error (SE), degrees of freedom, t-ratio, and p-values are given.

| Contrast | Estimate | SE | df | t.ratio | p.value |
| --- | --- | --- | --- | --- | --- |
| degraded - healthy | -4 | 0.75 | 39 | -5.27 | 1.56E-05 |
| degraded – outplant | -2.25 | 0.75 | 39 | -2.96 | 0.013 |
| healthy - outplant | 1.75 | 0.75 | 39 | 2.30 | 0.066 |

**Table S8 Output from a generalised linear model (family = Gamma) investigating the effect of site on sound richness.** Response refers to estimated marginal means predictions back transformed to the response scale. Standard error (SE), degrees of freedom (df) and confidence levels are given. The model explains a significant proportion of the variance in sound richness (null deviance = 10.1461 on 47 degrees of freedom, residual deviance = 7.1065 on 42 degrees of freedom).

| Site | Response | SE | df | lower.CL | upper.CL |
| --- | --- | --- | --- | --- | --- |
| degraded | 2.86 | 0.26 | 42 | 2.41 | 3.53 |
| healthy | 4.16 | 0.38 | 42 | 3.50 | 5.12 |
| outplant | 3.66 | 0.34 | 42 | 3.08 | 4.51 |

**Table S9 Post-hoc comparisons of model estimates of sound richness provided in Table S8, adjustment method was Tukey.** Model estimates, standard error (SE), degrees of freedom, t-ratio, and p-values are given.

| Contrast | Estimate | SE | df | t.ratio | p.value |
| --- | --- | --- | --- | --- | --- |
| degraded - healthy | 0.10 | 0.038 | 42 | 2.79 | 0.020 |
| degraded - outplant | 0.075 | 0.040 | 42 | 1.86 | 0.16 |
| healthy - outplant | -0.032 | 0.032 | 42 | -0.99 | 0.58 |

**Figure S1 Visual inspection of spectrograms for anthropogenic sound.** The upper image contains diver breathing, visible in repeated vertical stripes. The lower image contains boat noise, in the far right of the spectrogram where the colours turn to bright white. (Screenshots taken from Audacity ® sound analysis program (https://www.audacityteam.org/)).

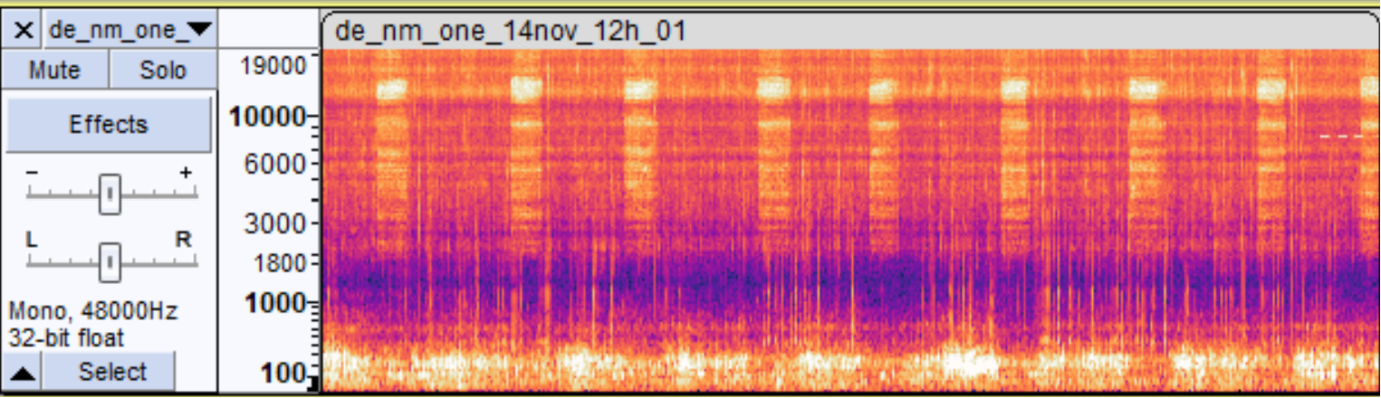

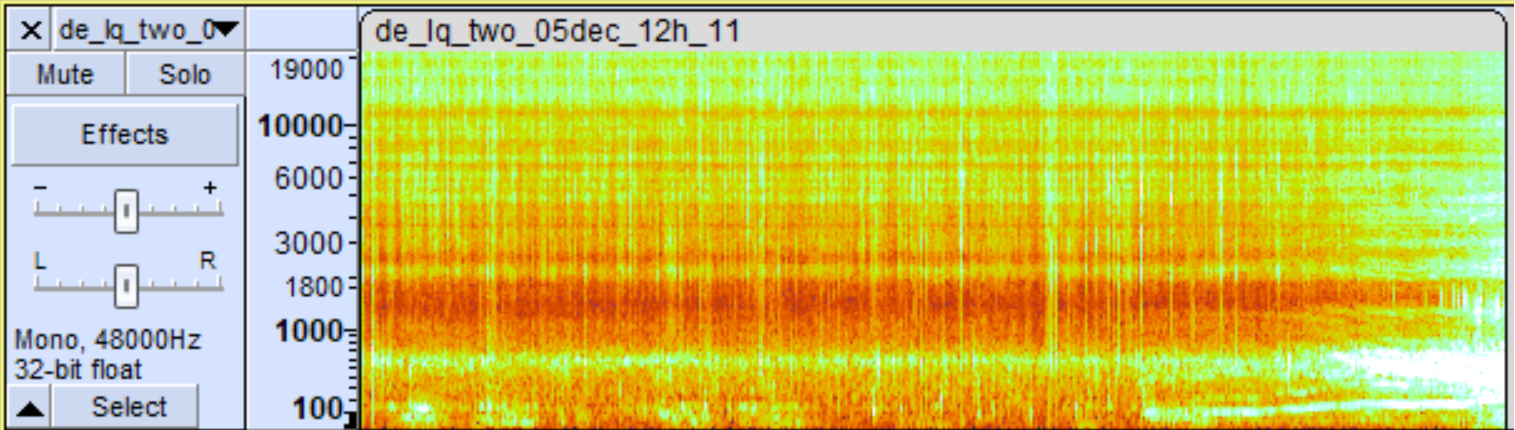

**Figure S2 UMAP visualisation of soundscape clustering by time of day.** UMAP Dimensions 1 and 2 refer to the axes of greatest variation. Each point represents a 5-second audio sample from a recording. Colours refer to the time of day, where 18h = sunset, 24h = midnight, 06h = sunrise, and 12h = midday.

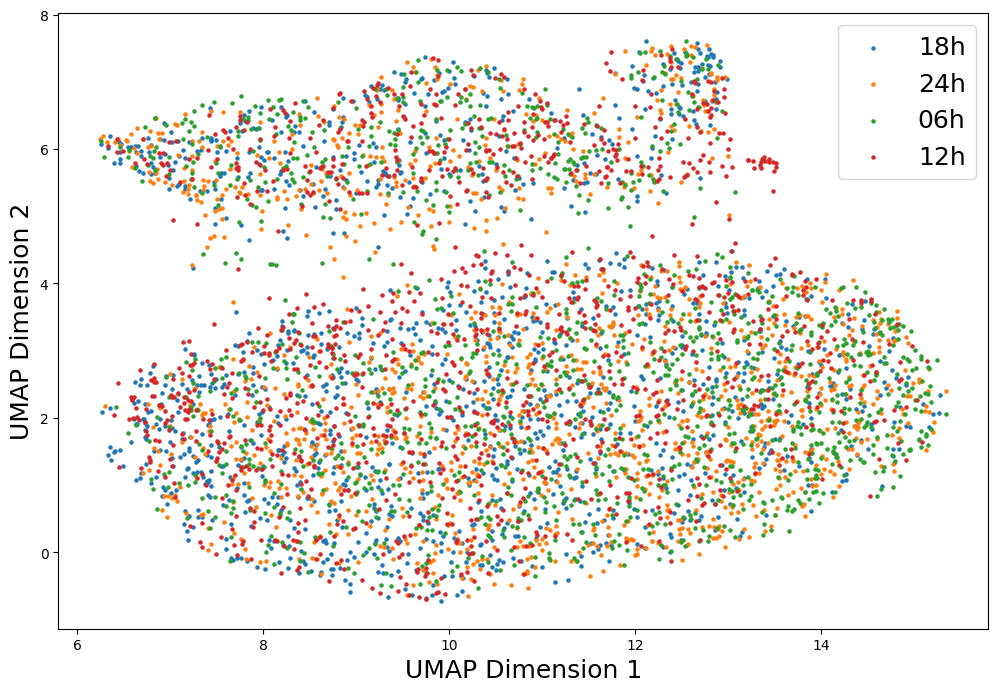

**Figure S3 UMAP visualisation of soundscape clustering, with unfiltered audio recordings as input (entire bandwidth).** UMAP Dimensions 1 and 2 refer to the axes of greatest variation. Each point represents a 5-second audio sample from a recording.

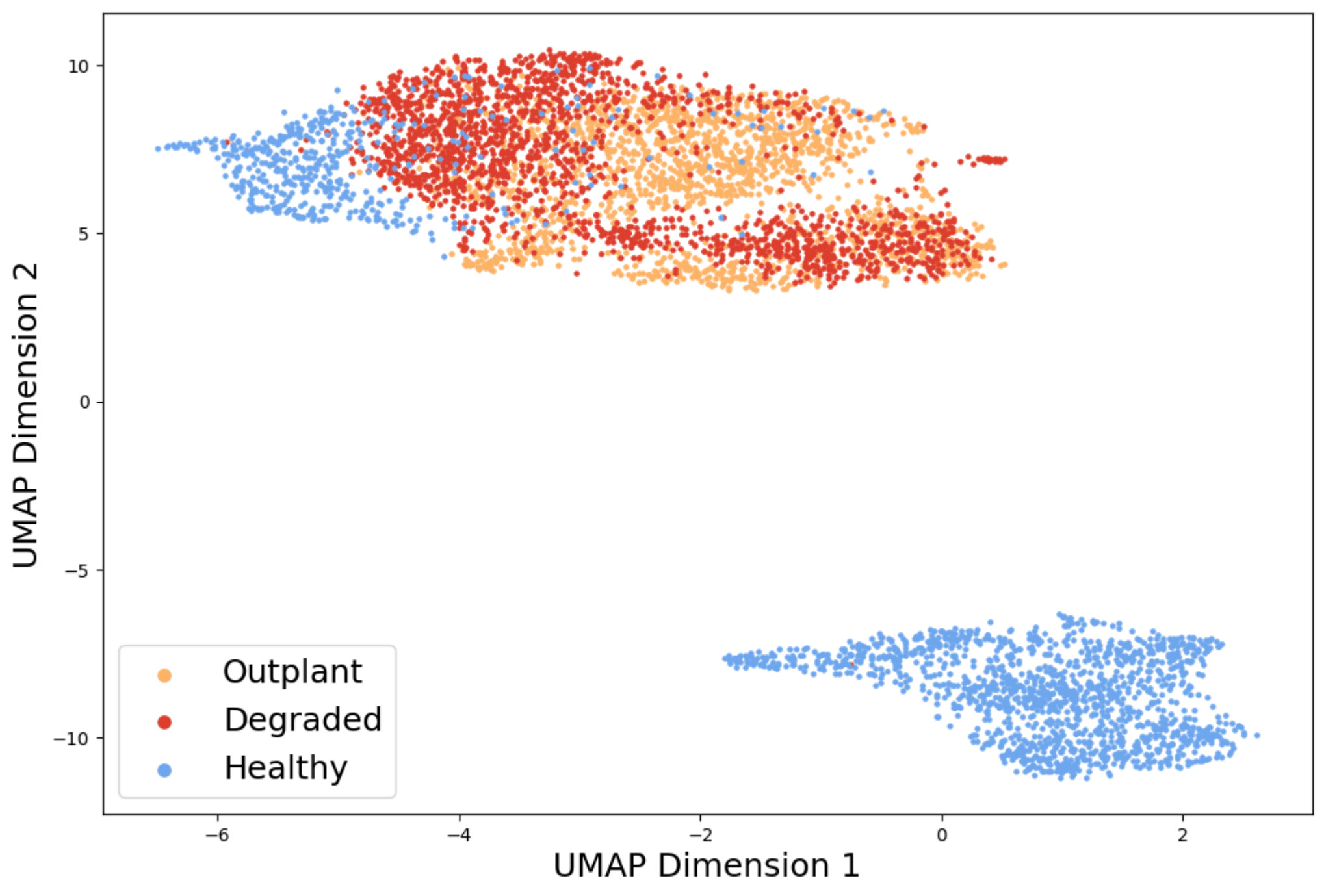

**Figure S4 UMAP visualisation of soundscape clustering, with recordings included in the manual analysis outlined in black.** UMAP Dimensions 1 and 2 refer to the axes of greatest variation. Each point represents a 5-second audio sample from a recording.
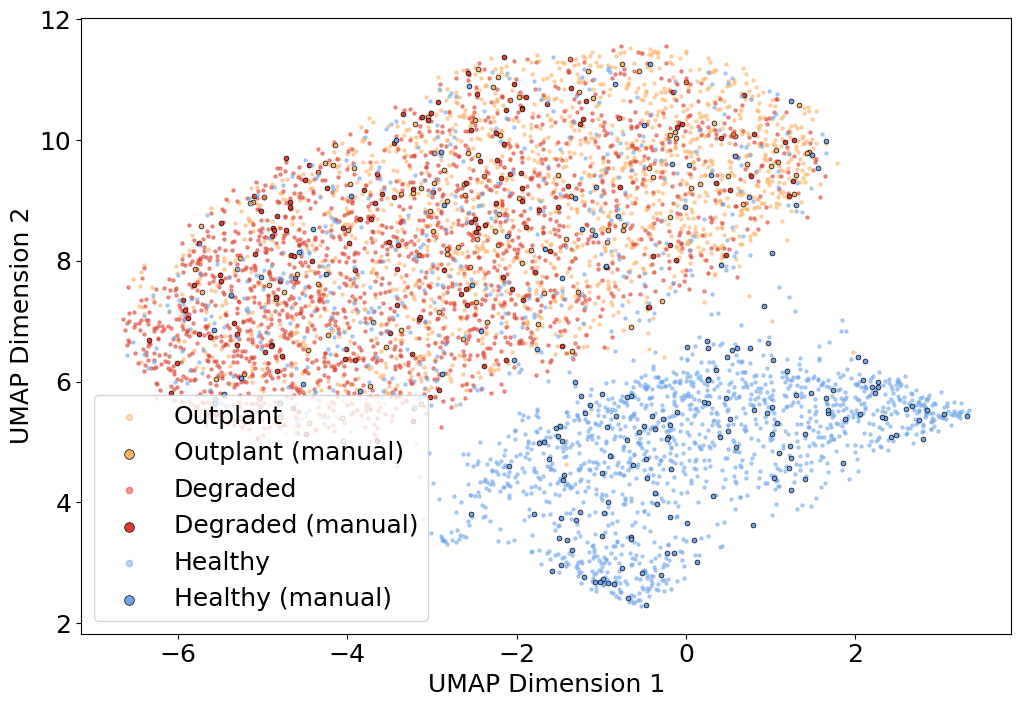

**Figure S5 UMAP exclusively with all the recordings that were manually analysed.** UMAP Dimensions 1 and 2 refer to the axes of greatest variation. Each point represents a 5-second audio sample from a recording.

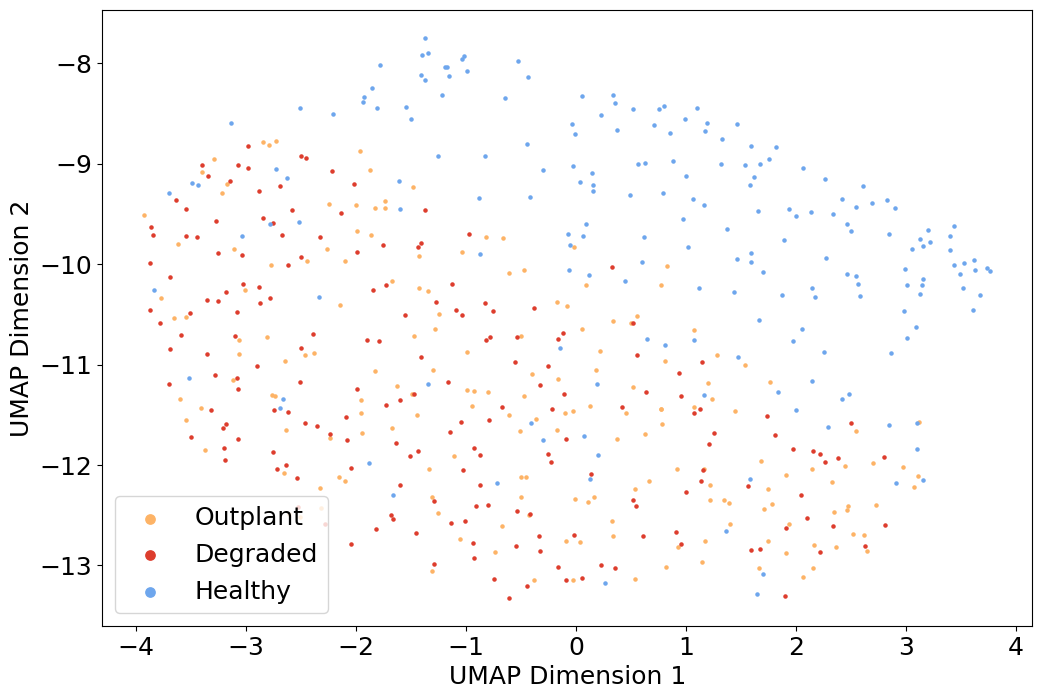
